# An *arginase 2* promoter transgenic illuminates anti-inflammatory signalling in zebrafish

**DOI:** 10.1101/2022.02.14.480079

**Authors:** Ffion R. Hammond, Amy Lewis, Holly E. Anderson, Lewis G. Williams, Annemarie H. Meijer, Geert F. Wiegertjes, Philip M. Elks

**Affiliations:** The Bateson Centre, Department of Infection and Immunity and Cardiovascular Disease, University of Sheffield, Western Bank, Sheffield, UK; Institute of Biology Leiden, Leiden University, Einsteinweg 55, 2333 CC, Leiden, The Netherlands; Aquaculture and Fisheries Group, Department of Animal Sciences, Wageningen University & Research, 6700 AH Wageningen, The Netherlands

## Abstract

The innate immune response to inflammatory stimuli must be finely balanced to produce an appropriate pro-inflammatory response while allowing a subsequent return to homeostasis. In recent years, *in vivo* transgenic zebrafish models have shed light on the temporal regulation of the pro-inflammatory innate response to immune challenges. However, until now, there have been no zebrafish transgenic models of anti-inflammatory signalling. We compared existing expression data of arginase genes in zebrafish neutrophils and macrophages, strong candidates for an anti-inflammatory marker, and identified that *arginase 2* is the most highly expressed Arginase in zebrafish immune cells. We developed an *arginase 2* (*arg2*) bacterial artificial chromosome (BAC) transgenic line, *TgBAC(arg2:eGFP)sh571*, driving GFP expression under the control of the *arg2* promoter. We show that, under resting conditions, *arg2:GFP* is expressed in ionocytes, matching the *in situ* hybridisation pattern. Upon immune challenge by injury, bacterial and fungal insults, *arg2:GFP* is predominantly expressed in neutrophils at early timepoints post-insult. Later in infections, *arg2:GFP* is expressed in cells associated with foci of infection (including neutrophils and macrophages), alongside liver expression. Our data indicate that *arginase 2* is predominantly expressed in neutrophils after immune challenge and suggest that anti-inflammatory signals coincide with pro-inflammatory signals during early wound and infection responses.

## Introduction

Initial immune responses to inflammatory or infection stimuli are mediated by innate immune cells, of which leukocytes are major players. Two important leukocyte cell types, neutrophils and macrophages, work together to neutralise invading threats whilst promoting tissue healing and restoration of homeostasis. Neutrophils and macrophages have evolved together and co-exist in invertebrate and vertebrate species [1]. Neutrophils are often observed to be the first immune cell type to respond to immune challenge, rapidly migrating towards the stimuli and becoming activated to a pro-inflammatory and antimicrobial state, clearing damaged cells and invading pathogens. Macrophages also rapidly respond to challenge, with pro-inflammatory phenotypes emerging soon after immune challenge aiding microbial clearance (often termed M1 or classical activation). Dogmatically, neutrophils are considered a blunt first line of defense that once activated remain so until cleared from tissues, either by apoptosis and subsequent efferocytosis by macrophages or by migration away [2–4]. Persistence of neutrophils during disease can cause considerable bystander damage to healthy surrounding tissues driving chronic pathologies. Macrophage phenotypes, on the other hand, are well-characterised as being plastic throughout the pathogenesis of inflammation and infections in a process termed macrophage polarisation. Pro-inflammatory macrophage phenotypes are followed by anti-inflammatory phenotypes that promote healing and restoration of homeostasis (often termed M2 or alternatively activated). Many of the above observations come from *in vitro* approaches. *In vivo* exploration indicates that macrophage polarisation is not binary but rather a spectrum of phenotypes and behaviours, and that the neutrophil response is more plastic than previously thought, with emerging roles for neutrophil subsets in tissue protection and healing [5–7].

Both neutrophils and macrophages are activated by danger and pathogen associated molecular patterns (DAMPs and PAMPs), which trigger the production of pro-inflammatory signals (e.g. IL-1β and TNF-α) after immune challenge [8,9]. These activate the production of antimicrobial molecules, including reactive nitrogen species (RNS), via the enzyme inducible nitric oxide synthase (iNOS) [10]. Immunomodulatory signals are required to limit and eventually turn-off the pro-inflammatory response in order for tissues to regain homeostasis. The best characterised of these anti-inflammatory signaling molecules are cytokines including the interleukins (IL), IL-4, IL-13 and IL-10, alongside proteins that dampen the pro-inflammatory response, including Arginase [11]. Despite their classification as anti-inflammatory, these signals have also been identified as being upregulated in some pro-inflammatory situations [12]. The production of antimicrobial RNS is negatively regulated by the enzyme Arginase. Arginase competes with iNOS for a shared substrate, L-arginine, a limited resource within the cell. The immunomodulatory properties of Arginase extend beyond the regulation of RNS production. Arginase 2 is essential for IL-10 mediated downregulation of pro-inflammatory factors making it a key anti-inflammatory enzyme [13]. Arginase and iNOS have been well-characterised in murine models as being strongly expressed in macrophage subtypes, with Arginase being a wound-healing/anti-inflammatory macrophage marker and iNOS a pro-inflammatory macrophage marker [14]. In human macrophages the distinction between iNOS and Arginase expressing macrophage subtypes is less well-defined, due, in part, to lower macrophage RNS in humans compared to mice. Interestingly, human neutrophils constitutively express Arginase with levels increasing after infection *in vitro*, but its roles are poorly understood *in vivo* [15]. Considering that Arginase is not expressed highly by murine neutrophils [16,17], alternative *in vivo* models are required to investigate the roles of Arginase in immunomodulation and human disease.

The zebrafish has become a powerful model organism to determine the molecular mediators of immunity against injury and pathogenic challenges [18,19]. Advantages of the zebrafish model include transparent larvae combined with fluorescent transgenic lines, allowing detailed microscopy in an intact organism *in vivo*. A key turning-point in zebrafish immunity research came with the development of transgenic lines marking neutrophil and macrophage cell populations. Initially, transgenic lines were developed that labelled whole populations of immune cells in zebrafish larvae, e.g., *TgBAC(mpx:GFP)i114* labelling the total neutrophil population and *Tg(mpeg1:GFP)* labelling the total macrophage population [20,21]. More recently, transgenic lines of important pro-inflammatory cytokines have become available (e.g., *TgBAC(tnfa:GFP)pd1028* and *TgBAC(il-1beta)sh445*)) allowing in-depth analysis of the cells producing these important signals following a variety of immune challenges *in vivo* [22,23]. These transgenic lines utilised bacterial artificial chromosome (BAC) technology, that allows tens of kilobases of promoter region to be used to drive the fluorescent protein, ensuring that the expression of the transgene recapitulates endogenous expression patterns as closely as possible. A key gap in this zebrafish “transgenic toolbox” is an anti-inflammatory fluorescent transgenic line. We therefore set out to develop an *arginase 2* promoter driven BAC transgenic line to understand the *arginase* response to immune challenge *in vivo*.

Here, we compared existing expression data of *arginase* genes in zebrafish neutrophils and macrophages and show that *arg2* is the most highly expressed *arginase* in zebrafish immune cells. We developed a new BAC transgenic line that drives GFP under the control of the *arginase 2* promoter and demonstrate that the *arg2:GFP* transgene is expressed in ionocytes, a population of skin resident cells, in resting conditions. Following a range of immune challenges, including tailfin transection (sterile injury), *Mycobacterium marinum* (bacterial), and *Cryptococcus neoformans*/*Candida albicans* (fungal) infections, *arg2:GFP* was predominantly upregulated in neutrophils. We identify a small population of macrophages that express *arginase 2* after injury and infection suggesting the presence of anti-inflammatory macrophages in zebrafish. The *arg2:GFP* transgenic line has the potential to uncover new mechanisms behind innate immune regulation during *in vivo* immune challenge.

## Materials and Methods

### Ethics statement and zebrafish

All zebrafish were raised in the Biological Services Unit (BSU) aquarium (University of Sheffield, UK) and maintained according to standard protocols (zfin.org) in Home Office approved facilities. All procedures were performed on embryos less than 5.2 dpf (days post fertilisation) which were therefore outside of the Animals (Scientific Procedures) Act, to standards set by the UK Home Office. Adult fish were maintained at 28°C with a 14/10 hour light/dark cycle. To investigate expression in immune cells the *TgBAC(arg2:eGFP)sh571* (hereon in termed *arg2:GFP*) reporter line was crossed with the macrophage reporter *Tg(mpeg1:mCherryCAAX)sh378* [24] (hereon in termed *mpeg:mCherry*) and the neutrophil reporter *Tg(lyz:nfsB*.*mCherry)sh260* [25] (hereon in termed *lyz:mCherry*) to generate embryos for experiments.

### Generation of *TgBAC(arg2:eGFP)sh571* transgenic zebrafish

An eGFP SV40 polyadenylation cassette was inserted at the *arg2* ATG start site of the zebrafish bacterial artificial chromosome (BAC) CH-211-12d10 using established protocols [21]. Inverted Tol2 elements were inserted into the chloramphenicol coding sequence and the resulting modified BAC containing 115130 base pairs of the *arg2* promoter region was used. We identified two founder zebrafish (allele codes sh571 and sh572) and raised colonies. The embryos of both alleles had the same GFP expression pattern. The data generated in this manuscript is from the *TgBAC(arg2:eGFP)sh571* transgenic line (hereon in termed *arg2:GFP*) as this line had a higher fecundity than *TgBAC(arg2:eGFP)sh572*.

### Tailfin transection

To induce an inflammatory stimulus, 2- or 3-days post fertilisation (dpf) zebrafish were anaesthetised in 0.168 mg/ml Tricaine (Sigma-Aldrich) in E3 media and visualised under a dissecting microscope. Using a scalpel blade (5mm depth, WPI) the tailfin was transected after the circulatory loop as previously described ensuring the circulation remained intact [26].

### Pathogen strains and culture

Bacterial infection experiments were performed using *Mycobacterium marinum* strain M (ATCC #BAA-535), containing the pSMT3-Crimson vector, with liquid cultures prepared from bacterial plates [27]. Liquid cultures were washed and prepared in 2% polyvinylpyrrolidone40 (PVP40) solution as previously described for injection [28,29]. Injection inoculum was prepared to 100 colony forming units (cfu)/nl for all *Mm* experiments, which was injected into the circulation at 30hpf via the caudal vein.

Fungal infection experiments were performed using the *Candida albicans* strain TT21*-mCherry* [30]. Overnight liquid cultures were grown from fungal plates, then prepared for injection as previously described [30]. Cultures were counted using a hemocytometer and prepared in 10% PVP40 for 200cfu/nl injection dose which was injected into the circulation at 30hpf via the caudal vein.

Fungal infection experiments were also performed using the *Cryptococcus neoformans* strain Kn99*-mCherry* [31]. Cryptococcal culture was performed as previously described [24] and after counting on a hemocytometer, Kn99 was prepared in 10% PVP40 for 200cfu/nl injection dose which was injected into the circulation at 1-2dpf.

### Microinjection of zebrafish larvae

Prior to injection, zebrafish were anaesthetised in 0.168 mg/ml Tricaine (MS-222, Sigma-Aldrich) in E3 media and transferred onto 1% agarose in E3+methylene blue plates, removing excess media. All pathogens were injected into the circulation to create systemic infection, using a microinjection rig (WPI) attached to a dissecting microscope. A 10mm graticule was used to measure 1nl droplets for consistency, and droplets were tested every 5-10 fish and recalibrated if necessary. Final injection volume of 1nl was injected to produce doses calculated for each pathogen. After injection, zebrafish were transferred to fresh E3 media for recovery and maintained at 28°C.

### Whole mount *in situ* hybridisation

RNA probes for zebrafish arginase type II (*arg2*, ENSDARG00000039269. Plasmid obtained from Source Bioscience) were designed and synthesised after cloning into the pCR™Blunt II-TOPO® vector according to manufacturer’s instructions (ThermoFisher Scientific). Plasmids were linearised and probes synthesised according to DIG RNA Labeling Kit (SP6/T7) (Roche). Zebrafish larvae were fixed in 4% paraformaldehyde solution (PFA, ThermoFisher Scientific) overnight at 4°C. Whole mount *in situ* hybridisation was performed as previously described [32].

### Confocal microscopy

Control, tailfin transected, and infected larvae were imaged using a Leica DMi8 SPE-TCS microscope using a HCX PL APO 40x/1,10 water immersion lens. For confocal microscopy larvae were anaesthetised in 0.168 mg/ml Tricaine and mounted in 1% low melting agarose (Sigma) containing 0.168 mg/ml tricaine (Sigma) in 15μ-Slide 4 well glass bottom slides (Ibidi).

### Stereo microscopy

Zebrafish larvae were anaesthetised in 0.168 mg/ml Tricaine and transferred to a 50mm glass bottomed FluoroDish™ (Ibidi). Zebrafish were imaged using a Leica DMi8 SPE-TCS microscope fitted with a Hamamatsu ORCA Flash 4.0 camera attachment using a HC FL PLAN 2.5x/0.07 and HC PLAN APO 20x/0.70 dry lens. Both transgenic zebrafish and whole mount *in situ* staining was imaged using a Leica MZ10F stereo microscope fitted with a GXCAM-U3 series 5MP camera (GT Vision).

### Lightsheet microscopy

2 and 3dpf larvae were imaged using a Zeiss Z1 lightsheet microscope with Plan-Apochromat 20x/1.0 Corr nd=1.38 objective, dual-side illumination with online fusion and activated Pivot Scan at 28°C chamber incubation. Zebrafish were anaesthetised in 0.168 mg/ml Tricaine and mounted vertically in 1% low melting agarose (Sigma) in a glass capillary. Images were obtained using 16bit image depth, 1400×1400 pixel field of view and GFP visualised with 488nm laser at 16% power, 49.94ms exposure and user-defined z-stack depth (400-600 slices, 0.641μm slices).

### Data analysis

Microscopy data was analysed using Leica LASX (Leica Microsystems) and Image J software. Graphs were generated using Prism 8.0, GraphPad Software (San Diego, CA, USA).

## Results

### *TgBAC(arg2:GFP)sh571* is expressed in ionocytes in resting conditions

There are two isozymes of Arginase in most mammals and fish, Arginase 1 (ARG1) and Arginase 2 (ARG2). In mice, ARG1 and ARG2 are both expressed by macrophages, however it is ARG1 that is the most widely studied in macrophage polarisation, with increased expression of cytosolic ARG1 protein depleting intracellular stores of arginine [33]. In fish, Arginases have been studied in common carp (*Cyprinus carpio*), a species phylogenetically close to zebrafish, where *arg2* is the most highly expressed of the Arginase family in immune cells [11,34]. Head-kidney derived macrophages of carp which have been polarised towards anti-inflammatory phenotypes using cAMP have a 16-fold upregulation of *arg2* expression [11,35].

Zebrafish have orthologues of mammalian Arginase 1 and Arginase 2, which were compared using existing transcriptomics datasets to identify innate immune cell expression. FACS purified neutrophils (*Tg(mpx:GFP)i114*) and macrophages (*Tg(mpeg1:Gal4-VP16)gl24/(UAS-E1b:Kaede)s1999t*) from unchallenged 5dpf larvae both expressed *arginase2* (*arg2*), while *arginase 1* (*arg1*) was not expressed at detectable levels (Supplemental Figure 1A-B, using raw data from [36]). *arg2* expression was found at high levels in the bulk, non-immune cell, population alongside being expressed in the immune cell populations (Supplemental Figure 1C-D, data from [36]). *arg2* expression was approximately 1.5-fold higher in neutrophils than macrophages in unchallenged zebrafish larvae (Supplemental Figure 1C-D, using raw data from [36]). In unchallenged adult zebrafish, single-cell RNAseq identified *arg2* expression in a population of neutrophils, a smaller population of monocytes/macrophages, with no detectable expression in other blood lineages (thrombocytes/erythrocytes) (Supplemental Figure 1E-F, data from The Zebrafish Blood Atlas, https://www.sanger.ac.uk/science/tools/basicz/basicz/, [37]). From the same dataset *arg1* was not detected in any immune cell lineage (Supplemental Figure 1E-F).

Based on the predominant expression of *arg2* in fish innate immune cells, we chose to develop an *arg2* transgenic zebrafish line to investigate its expression during immune challenge *in vivo*. We adopted a BAC transgenesis approach, using a BAC (CH-211-12d10) in which 11.5kb of the *arg2* promoter drives GFP expression, to generate two transgenic line alleles with the same expression (*TgBAC(arg2:eGFP)sh571* and *TgBAC(arg2:eGFP)sh572*). Due to the higher fecundity in the *sh571*, it was this line that was used in the following studies (hereon in termed *arg2:GFP*).

In unchallenged *arg2:GFP* larvae, the transgene was expressed in cuboidal-shaped cells in the skin, distributed over the yolk and caudal vein regions at 2 days post fertilisation (dpf, Figure 1A) and 3dpf (Figure 1B-C). A subset of ionocytes, cells in the skin responsible for the transport of sodium ions also known as H^+^-ATPase-rich cells (HRCs), have been shown to express high levels of *arginase 2* by *in situ* hybridisation in zebrafish [38–40]. *arg2:GFP* expression recapitulated the *arginase 2 in situ* hybridisation pattern, labelling ionocytes (Figure 1D-E). To determine whether any of the cells over the yolk area expressing *arg2:GFP* were leukocytes, we crossed the *arg2:GFP* line with a macrophage transgenic line (*Tg(mpeg1:mCherry)sh378*, hereon in termed *mpeg:mCherry*) and a neutrophil transgenic line (*Tg(lyz:nsfB*.*mCherry)sh260*, hereon in termed *lyz:mCherry*). Under resting conditions there was no overlap between *mpeg:mCherry* positive macrophages and *arg2:GFP* positive cells at 2dpf, 3dpf, 4dpf or 5dpf (Figure 1F-G). Nor was there overlap between *lyz:mCherry* positive neutrophils (Figure 1H). These data indicate that the *arg2:GFP* line is not expressed at detectable levels in immune cells in resting conditions at these developmental timepoints, matching *in situ* hybridisation data. The zebrafish *arg2:GFP* transgene and *in situ* hybridisation did not detect leukocyte *arginase 2* expression in resting conditions, however *arginase 2* levels were detected by RNAseq of FACS purified leukocyte populations (Supplemental Figure 1). This suggests that either the transgenic and *in situ* hybridisation techniques were not as sensitive as RNAseq, or that techniques to purify immune cells in RNAseq studies have led to upregulation of *arg2* not present in the intact zebrafish.

**Figure 1:**
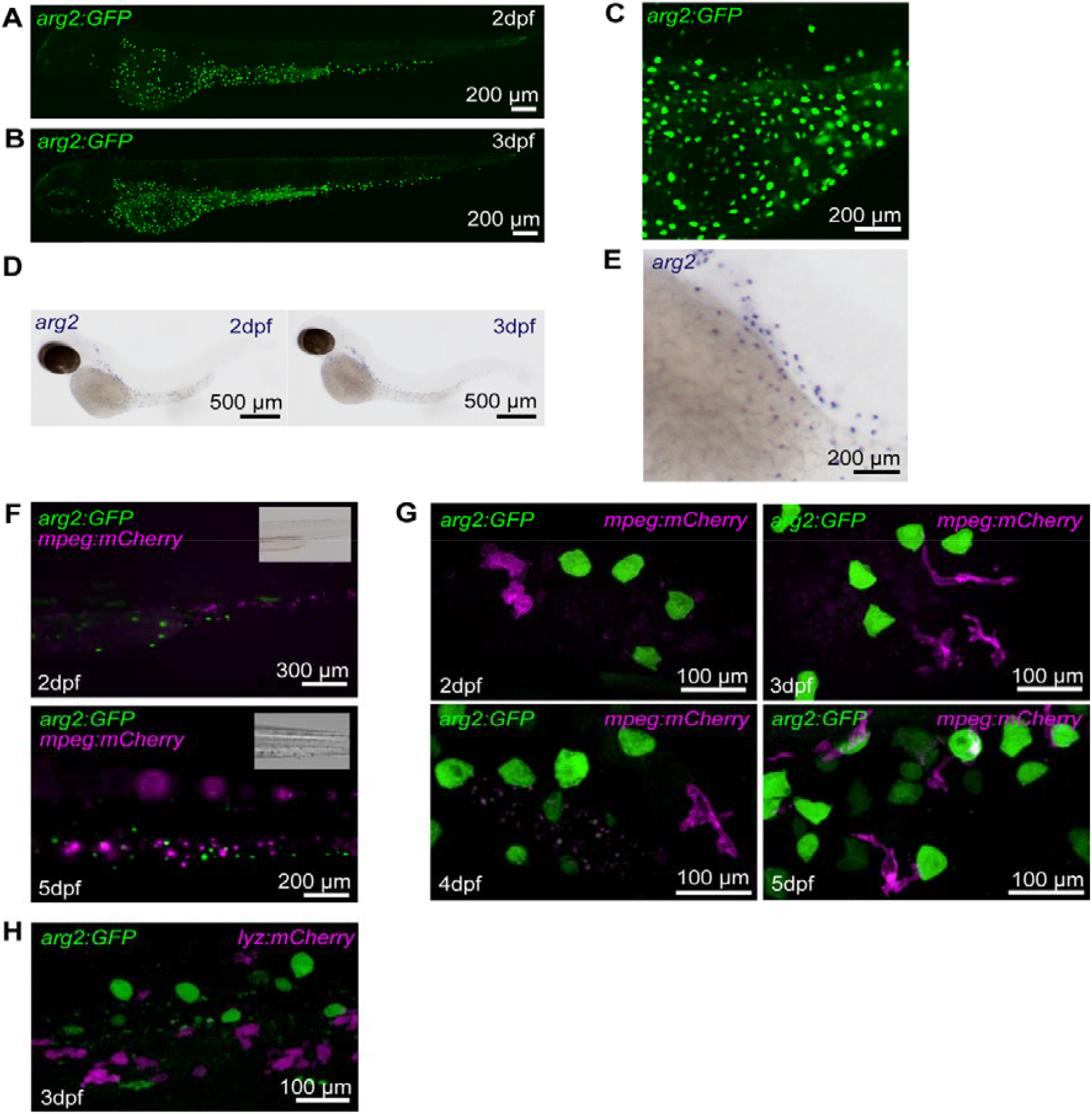
The *TgBAC(arg2:eGFP)sh571* line shows GFP expression in ionocytes but not in resting macrophages and neutrophils, recapitulating the *arginase 2 in situ* hybridisation expression pattern. (A-B) Lightsheet microscopy stereo micrographs of *TgBAC(arg2:eGFP)sh571* line shows ionocytes expression at 2 (A) and 3dpf (B). (C) Enlarged image of section over yolk of (B). (D) *arginase 2 in situ* hybridisation shows expression in cells over the yolk known as ionocytes at 2 and 3 dpf in unchallenged zebrafish. (E) Enlarged image of section over yolk of 3dpf larvae in (D). (F) Stereo fluorescence micrographs of *arg2:GFP* crossed to *Tg(mpeg1:mCherry)sh378* at 2dpf and 5dpf with no overlap of *arg2:GFP* expression in ionocytes with magenta macrophages. (G) Stereo fluorescence micrograph of *arg2:GFP* crossed to *Tg(mpeg1:mCherry)sh378* at 2dpf, 3dpf, 4dpf and 5dpf with no overlap of *arg2:GFP* expression in ionocytes with magenta macrophages. (H) Stereo fluorescence micrograph of *arg2:GFP* crossed to *Tg(lyz:nfsB*.*mCherry)sh260* at 3dpf with no overlap of *arg2:GFP* expression in ionocytes with magenta neutrophils.

### *arg2:GFP* is predominantly upregulated by neutrophils after tailfin transection

To assess whether *arg2:GFP* expression is upregulated in innate immune cells during an immune response, we challenged 2dpf larvae with a sterile tailfin wound. Using a tailfin nick model we identified that highly mobile immune cells migrating towards the wound expressed *arg2:GFP* within the first hour of timelapse microscopy (Figure 2A). We reasoned that these cells were neutrophils, due to their size and amoeboid shape alongside their rapid migration towards the wound. To investigate whether neutrophils expressed *arg2:GFP* after injury, we crossed the *arg2:GFP* line with the neutrophil *lyz:mCherry* transgenic line. Timelapse microscopy demonstrated that a subpopulation of neutrophils arriving at the tailfin transection wound were *arg2:GFP* positive by 1 hour post wound (hpw) (Figure 2B) with positive neutrophils at the site of injury present during the recruitment phase of inflammation (1-6hpw) while expression was not observable in neutrophils away from the wound. There was no expression of *arg2:GFP* observed in *mpeg:mCherry* positive macrophages in the recruitment phase of inflammation between 1-6hpw (Figure 2C).

**Figure 2:**
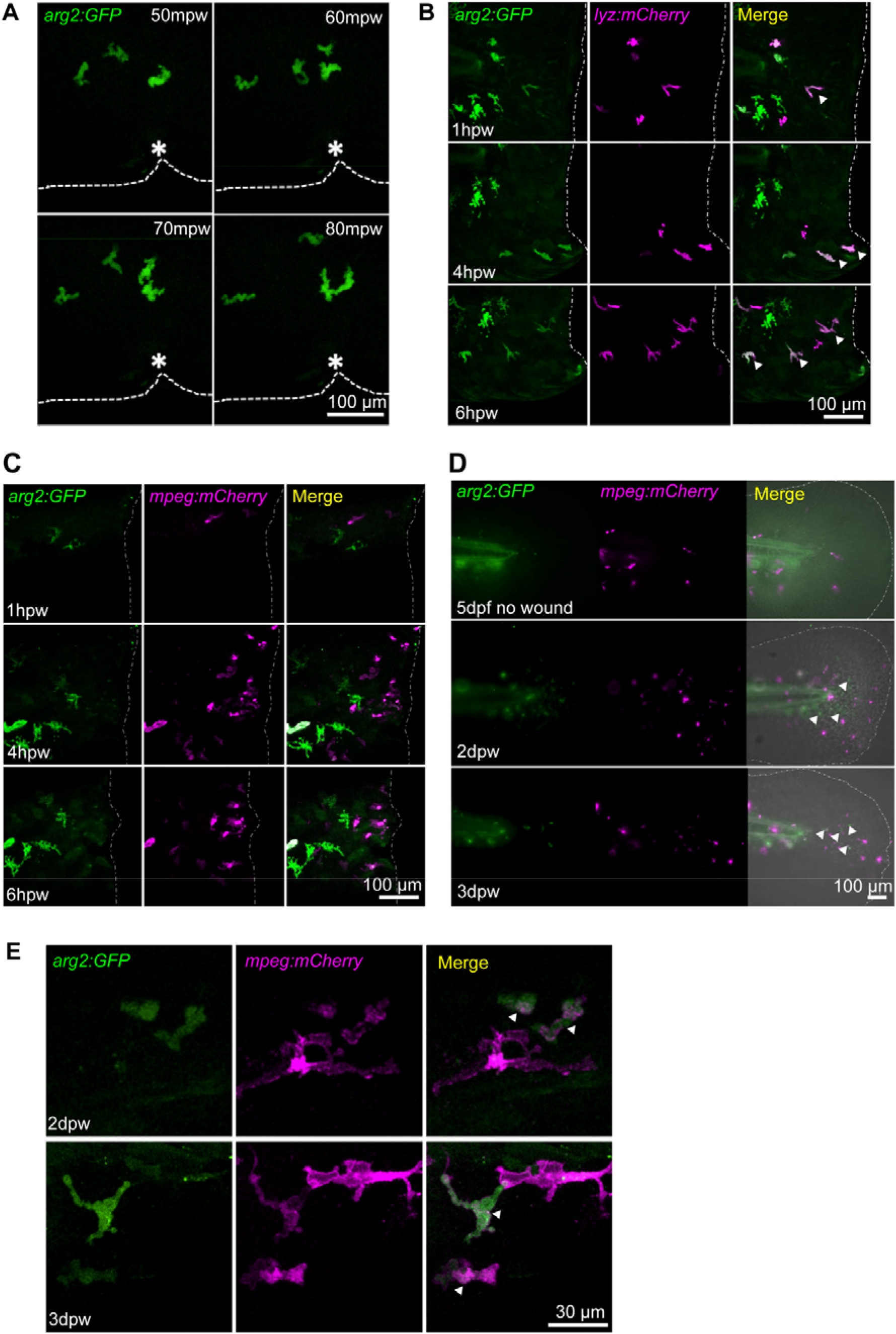
Neutrophils express *arg2:GFP* after wound challenge. (A) Fluorescence confocal timelapse micrographs of *TgBAC(arg2:eGFP)sh571* positive cells migrating towards a tailfin nick wound at early timepoints post injury. Dashed line indicates the edge of the fin and the asterisk marks the nick wound. (B) Fluorescence confocal micrographs of *arg2:GFP* crossed to *Tg(lyz:nfsB*.*mCherry)sh260* showing overlap of *arg2:GFP* with neutrophils at a tailfin wound (dashed line). Arrowheads indicate *arg2:GFP* positive neutrophils migrating at the wound. (C)Fluorescence confocal timelapse micrographs of *arg2:GFP* crossed to *Tg(mpeg1:mCherry)sh378* showing no overlap of macrophages with *arg2:GFP* expression early after injury (dashed line indicates the wound). (D) Fluorescence widefield micrographs of *arg2:GFP* crossed to *Tg(mpeg1:mCherry)sh378*. The upper panels show an uninjured tailfin with few macrophages and no immune cell *arg2:GFP* expression at 5dpf. The middle panels show an injured tailfin at 2dpw (4dpf) showing no overlap between *mpeg:mCherry* and *arg2:GFP*, but some unlabelled cells expressing *arg2:GFP* with amoeboid, immune cell shape (white arrowheads). Dotted line indicates the edge of the tailfin fold. The lower panels show an injured tailfin at 3dpw (5dpf) showing no overlap between *mpeg:mCherry* and *arg2:GFP*, but some unlabelled cells expressing *arg2:GFP* with amoeboid, immune cell shape (white arrowheads). Dotted line indicates the edge of the tailfin fold. (E) Fluorescence confocal micrographs of *arg2:GFP* crossed to *Tg(mpeg1:mCherry)sh378* at 2dpw (upper panels) and 3dpw (lower panels), showing examples of *mpeg:mCherry* positive *arg2:GFP* expressing cells in the proximity of the healing wound (arrowheads).

In order to observe potential anti-inflammatory macrophages, *mpeg:mCherry* larvae crossed into the *arg2:GFP* line were imaged at a later timepoint, 3dpw (5dpf), by which time the fin has partially regenerated. In uninjured larvae there were few *mpeg:mCherry* positive macrophages in the tailfin fold at 5dpf and these did not express *arg2:GFP* (Figure 2D, top panel). In tailfin transected larvae there were increased numbers of macrophages in the 2dpw and 3dpw healing/regenerating tailfin fold, but the majority of these were *arg2:GFP* negative, while *mpeg:mCherry* negative cells with the morphology of neutrophils did express *arg2:GFP* (Figure 2D, bottom two panels). Upon closer investigation using confocal microscopy, *mpeg:mCherry* positive cells expressing *arg2:GFP* were identified at the wound at 2dpw and 3dpw (Figure 2E), though these were greatly outnumbered by *arg2:GFP* negative macrophages. *arg2:GFP* positive macrophages were not observed in uninjured larvae. Together these data hint at the presence of *arginase 2* expressing anti-inflammatory macrophages during tailfin regeneration.

### Infection challenge upregulates neutrophil *arg2:GFP* expression

To investigate the expression of *arg2* after bacterial infection, *arg2:GFP* embryos were injected with *Mycobacterium marinum* (Mm) at 1dpf. We assessed the early response to Mm infection at 1dpi and found that a subpopulation of neutrophils express *arg2:GFP* early in infection (Figure 3A). Neutrophil *arg2:GFP* expression was not observed in mock-infected (PVP) controls (Figure 3B) with ionocyte expression confirming *arg2:GFP* positive larvae. *arg2:GFP* positive neutrophils were present in the vicinity of Mm infection (Figure 3A), with both infected and uninfected neutrophils expressing *arg2:GFP* (Figure 3C). A subset of neutrophils had *arg2:GFP* expression, with 31.7% (of n=123 neutrophils) of *lyz:mCherry* positive neutrophils in the region of infection expressing GFP, suggestive of differential immune responses between individual neutrophils (Figure 3D-E). At the same 1dpi timepoint, 6.1% (of n=98 macrophages) of *mpeg:mCherry* positive macrophages were positive for *arg2:GFP* (Figure 3E-G).

**Figure 3:**
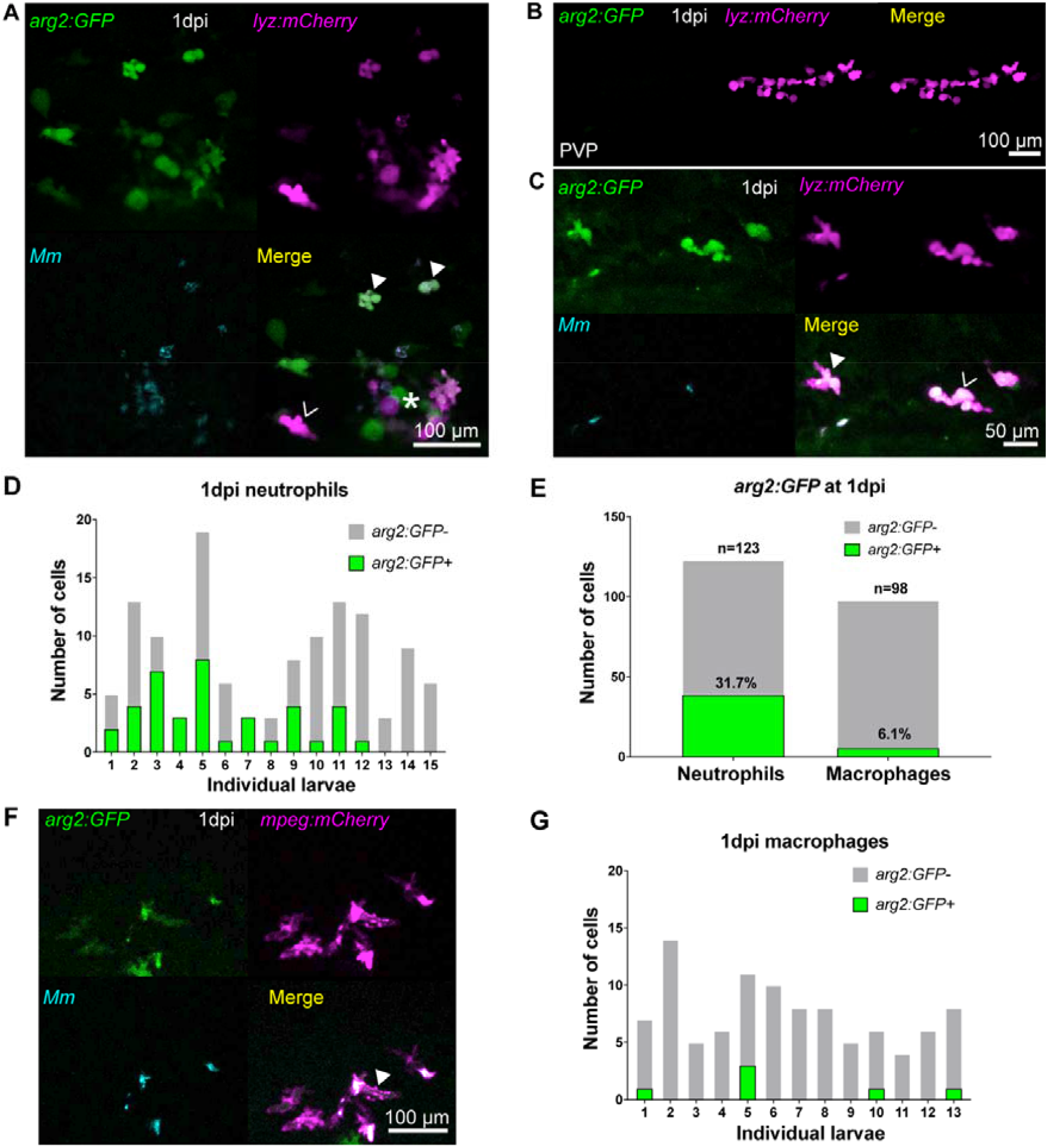
Neutrophils are the predominant immune cell that express *arg2:GFP* post Mm challenge. (A) Fluorescence confocal micrographs of *TgBAC(arg2:GFP)sh571* crossed to *Tg(lyz:nfsB*.*mCherry)sh260* after Mm infection at 1dpi showing GFP positive neutrophils (filled arrowheads) and GFP negative neutrophils (hollow arrowheads), around an area of high infection (asterisk). (B) Fluorescence confocal micrographs of 1dpi *arg2:GFP* crossed to *lyz:mCherry* after PVP control injection. (C) Fluorescence confocal micrographs of *arg2:GFP* crossed to *lyz:mCherry* after Mm infection at 1dpi showing that both infected (filled arrowhead) and non-infected (hollow arrowhead) neutrophils can express *arg2:GFP*. (D) Graph showing the number of *arg2:GFP* positive and negative neutrophils in a 40x region of interest in the caudal vein region that contained Mm bacteria, post infection, in individual larvae. (E) Graph showing the percentage of *arg2:GFP* positive and negative neutrophils and macrophages in a 40x region of interest around the infected caudal vein region. Data shown are n = 98-123 cells accumulated over 3 independent experiments. (F) Fluorescence confocal micrographs of *arg2:GFP* crossed to *Tg(mpeg1:mCherry)sh378* after Mm infection at 1dpi showing a single GFP positive macrophage (filled arrowhead) in this field of view. One of 6 instances observed. (G) Graph showing the number of *arg2:GFP* positive and negative macrophages in a 40x region of interest in the caudal vein region that contained Mm bacteria post infection in individual larvae.

Fungal infections have been shown to modulate host arginine metabolism via Arginase [41], therefore we assessed *arg2:GFP* expression in two well-characterised fungal zebrafish infection models - *Candida albicans* [42] and *Cryptococcus neoformans* [24]. In both fungal infections, *arg2:GFP* was observed in a subpopulation of *lyz:mCherry* positive neutrophils at 1dpi (Figure 4A-B). As with Mm infection, subsets of neutrophils both with or without internalised pathogen were *arg2:GFP* positive in *Candida albicans* infection (Figure 4A). *arg2:GFP* positive *mpeg:mCherry* macrophages were also observed in *Cryptococcus neoformans* infection at 1dpi, however these were greatly outnumbered by negative macrophages and only 3 examples were identified (Supplemental Figure 2). In Cryptococcal infected larvae with a high fungal burden, *arg2:GFP* expression was observed in the liver (Figure 4C-D) at timepoints when this was not present in PVP injected larvae (Figure 4C). This was especially apparent in Cryptococcus infections at 2dpi (Figure 4D) and was confirmed by *in situ* hybridisation (Figure 4E). Arginase is a well characterised liver enzyme [43], yet in unchallenged embryos (Figure 1A-E) or PVP injected larvae (Figure 4C) there was no visible liver expression of *arg2:GFP*. Liver expression was also present after Mm infection, but was less pronounced (Supplemental Figure 3). Like Cryptococcal infected larvae this occurred in individuals highly infected with Mm, but was not observed till later in infection (at 4dpi), potentially reflecting the slower doubling time/pathogenesis of Mm compared to Cryptococcus.

**Figure 4:**
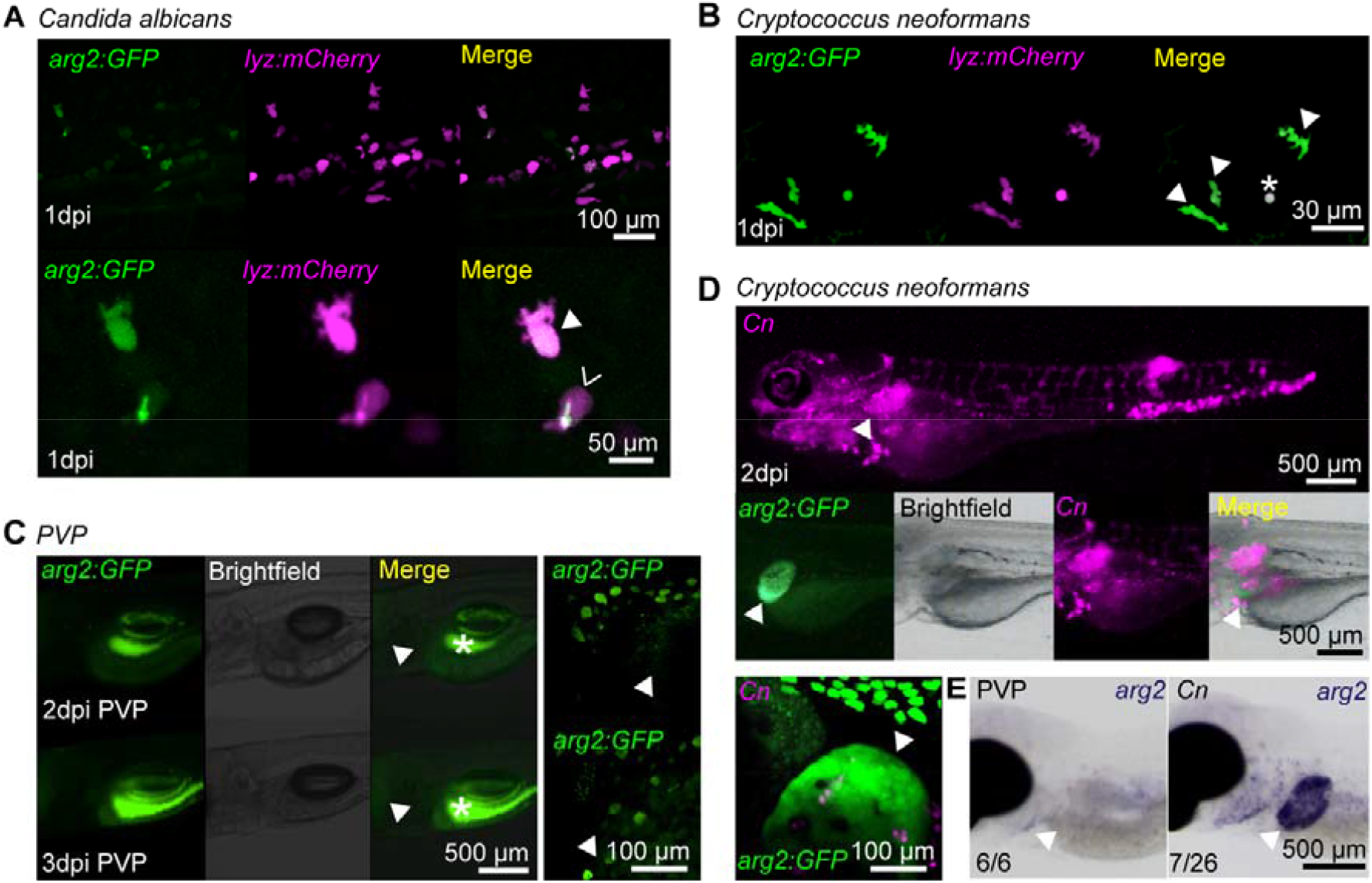
Fungal infections upregulate *arg2:GFP* in neutrophils and the liver. (A)Fluorescence confocal micrographs of *arg2:GFP* crossed to *lyz:mCherry* after *Candida albicans* infection at 1dpi showing GFP positive neutrophils (filled arrowhead) and GFP negative neutrophils (hollow arrowhead), with a zoomed in panel below showing a positive non-infected neutrophil and a GFP negative infected neutrophil (GFP in this cell is autofluorescence from the Candida, which in this instance has survived and formed a hyphae). (B) Fluorescence confocal micrographs of *arg2:GFP* crossed to *lyz:mCherry* after *Cryptococcus neoformans* infection at 1dpi showing GFP positive neutrophils (filled arrowheads). Asterisk indicates a Cryptococci that is auto fluorescent in both channels. (C) Brightfield and fluorescence micrographs of *arg2:GFP* larvae at 2dpi and 3dpi after PVP injection. The position of the *arg2:GFP* negative liver is shown by the arrowhead. The GFP channel is turned up highly enough to show gut fluorescence (asterisk) in order to show that the liver is *arg2:GFP* negative. (D) Brightfield and fluorescence micrographs of *arg2:GFP* larvae at 2dpi after *Cryptococcus neoformans* (Cn) infection showing *arg2:GFP* liver expression with heavy levels of infection. (E)Brightfield stereo micrographs of *arg2* wholemount *in situ* hybridisation after PVP or *Cryptococcus neoformans* (Cn) infection showing *arg2* liver expression (arrowhead) in an infected individual, not present in the PVP injected larvae.

### *arg2:GFP* is expressed in cells associated with developing granulomas

*arg2:GFP* expression was assessed at a later stage of Mm infection (4dpi), a stage at which innate immune granulomas are forming and when it is likely that leukocyte phenotypes are more diverse and polarised due to immune modulation by mycobacteria [44]. PVP control injected larvae had minimal *arg2:GFP* expression, apart from ionocyte expression and background green signal from the autofluorescence of pigment cells (Figure 5A). In Mm infected larvae, *arg2:GFP* was expressed in cells associated with developing granulomas at 4dpi (Figure 5B). The brightest population of these cells were *lyz:mCherry* positive, indicating a subset of *arg2:GFP* positive granuloma-associated neutrophils, representing 37.5% of neutrophils imaged in granulomas (Figure 5C-E). There were also granuloma-associated *arg2:GFP* expressing cells that were *lyz:mCherry* negative (Figure 5C), some containing phagocytosed bacteria (Figure 5C and F) and expressing macrophage *mpeg:mCherry* (Figure 5F). *mpeg:mCherry* positive macrophages had lower *arg2:GFP* expression in comparison to the *mpeg:mCherry* negative/*arg2:GFP* positive granuloma-associated cells (Figure 5F-H). *arg2:GFP* expression was observed in both infected and non-infected *mpeg:mCherry* positive macrophages (Figure 5F-H). Due to the dim and variable expression of both *arg2:GFP* and *mpeg:mCherry*, numbers of positive granuloma-associated macrophages were difficult to accurately quantitate, but *arg2:GFP* positive *mpeg:mCherry* macrophages were outnumbered by *arg2:GFP* negative *mpeg:mCherry* macrophages and *arg2:GFP* positive *lyz:mCherry* neutrophils in granulomas. Together, these data suggest that Mm granulomas have *arginase* expressing neutrophils and that other granuloma-associated cells, including potential anti-inflammatory macrophages, also express *arginase*.

**Figure 5:**
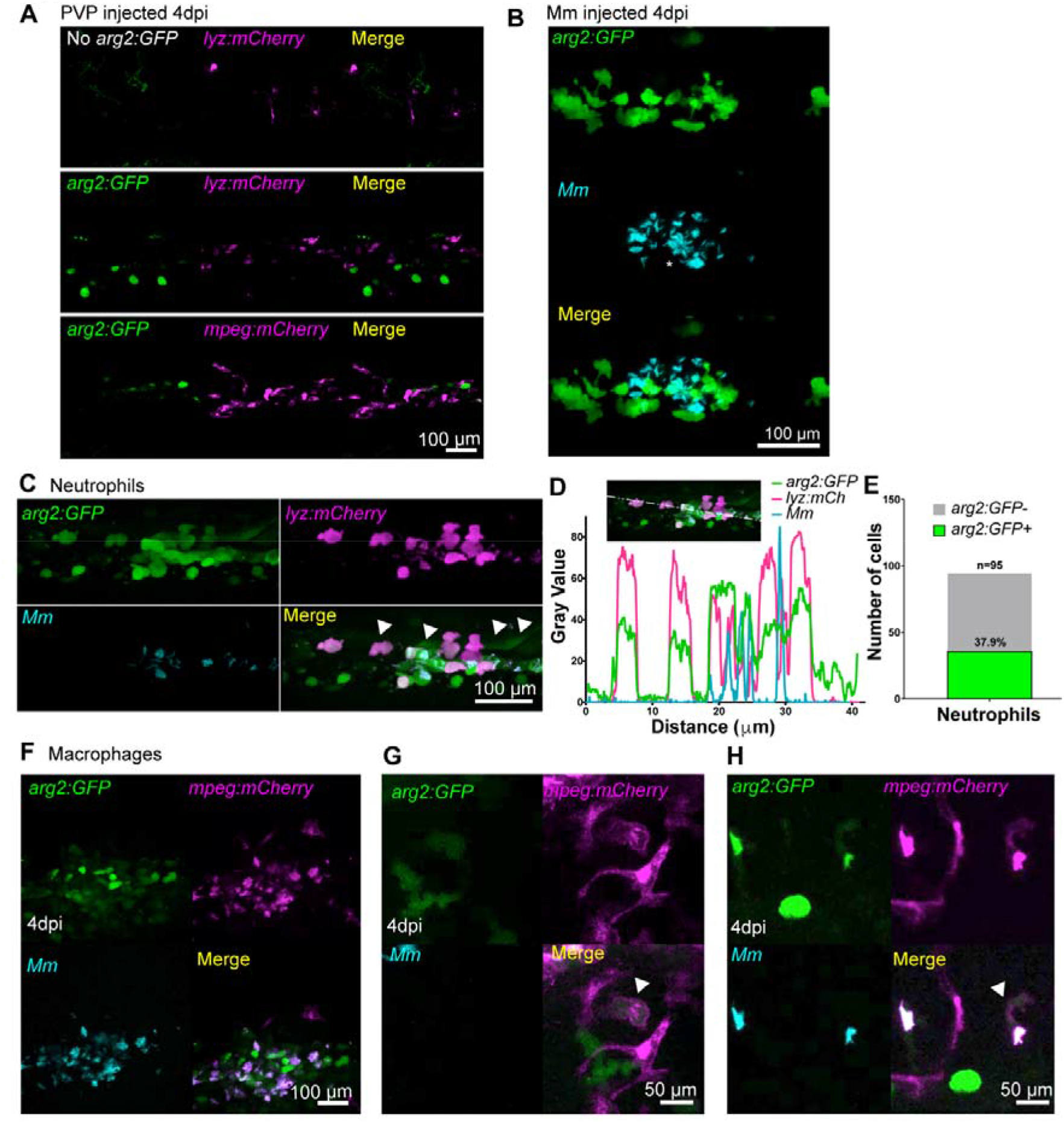
A subset of granuloma associated neutrophils and macrophages express *arg2:GFP*. (A) Fluorescence confocal micrographs of *TgBAC(arg2:GFP)sh571* crossed to *Tg(lyz:nfsB*.*mCherry)sh260* and *Tg(mpeg1:mCherry)sh378* after PVP control injection at 4dpi. Only ionocyte expression of *arg2:GFP* is present. (B) Fluorescence confocal micrographs of *TgBAC(arg2:GFP)sh571* after Mm infection at 4dpi showing granuloma associated *arg2:GFP* positive cells. (C) Fluorescence confocal micrographs of *arg2:GFP* crossed to *Tg(lyz:nfsB*.*mCherry)sh260* after Mm infection at 4dpi, showing neutrophils positive with *arg2:GFP* (filled arrowheads). (D) Line analysis of a cross section through a granuloma showing fluorescence values of *arg2:GFP, lyz:GFP* and *Mm*. (E) Graph showing the percentage of *arg2:GFP* positive and negative granuloma-associated neutrophils. Data shown are n = 95 cells accumulated over 3 independent experiments. (F) Fluorescence confocal micrographs of *TgBAC(arg2:GFP)sh571* crossed to *Tg(mpeg1:mCherry)sh378* after Mm infection at 4dpi. (G) Fluorescence confocal micrographs of *arg2:GFP* crossed to *mpeg:mCherry* after Mm infection at 4dpi showing a non-infected, *arg2:GFP* positive macrophage (filled arrowhead). (H) Fluorescence confocal micrographs of *arg2:GFP* crossed to *mpeg:mCherry* after Mm infection at 4dpi showing an infected, *arg2:GFP* positive macrophage (filled arrowhead).

## Discussion

Zebrafish transgenic lines have allowed *in vivo* exploration of the pro-inflammatory immune response (including *il-1β* and *tnfa*) to a variety of stimuli, however, until now, there has been a lack of similar tools for anti-inflammatory factors [22,23,45]. Here, we describe a new transgenic line for *arginase 2* and show that this transgene is upregulated shortly after immune challenge in neutrophils and a small subset of macrophages. These data correlate with a growing body of evidence that suggest anti-inflammatory factors are expressed early after immune challenge to suppress hyper-inflammation before a switch to an anti-inflammatory environment is required at later stages for tissue healing and restoration of homeostasis [46].

The *arg2:GFP* line shows that neutrophils are the predominant immune cell type that express *arginase 2* early after immune challenge in zebrafish. The observation of *arginase 2* expression in zebrafish neutrophils fits with our previous observations that zebrafish neutrophils are the primary innate immune cell that produce antimicrobial reactive nitrogen species (RNS) after Mm infection and suggests a neutrophil iNOS/Arginase balance [47]. Human neutrophils also produce high levels of RNS [12], differing from expression in mice where it is primarily macrophages that produce RNS/Arginase [12,16,48,49]. Human polymorphonuclear neutrophils (PMNs) produce high levels of Arginase at a transcript and protein level following immune challenge and in resting states, that may act as a negative regulator of the RNS response, as is the case in murine macrophages. [12,16,48,49].

The mechanisms balancing innate immune regulation during inflammatory and infection responses are not well understood *in vivo*. Neutrophil *arg2:GFP* was observed at timepoints that are considered to be pro-inflammatory stages of inflammation and infection. *arg2:GFP* expression was reminiscent of our previous observations using the pro-inflammatory *TgBAC(il-1 β:GFP)sh445* transgenic line, where neutrophil *il-1β* was found at 1hpw in the tailfin and 1dpi in Mm infections, the same timepoints that *arg2:GFP* was observed [23]. This suggests that anti-inflammatory *arginase* expression coincides with pro-inflammatory signals in neutrophils, evidence for a balanced response. The observation of immune cells producing anti-inflammatory signals upon immune challenge has been observed previously, but to date this has been mainly described in macrophages [46]. It is becoming clear that a balanced response to immune challenge, including both pro- and anti-inflammatory signals is beneficial in disease control [46]. In murine macrophages it has been demonstrated *in vitro* that Arginase 1 is produced soon after infection and can have immunomodulatory effects [16]. Our findings open up the possibility that a similar balance may exist in neutrophils and add to recent evidence suggesting that neutrophil phenotypes are more diverse and nuanced than previously appreciated [6,7].

Some pathogens have evolved to disrupt the pro- and anti-inflammatory response, keeping pro-inflammatory factors low and increasing anti-inflammatory signals to allow for immune cell evasion and survival. One such pathogen is *Mycobacterium tuberculosis* (Mtb), that suppresses an initial pro-inflammatory response, in part by upregulation of macrophage Arginase, to allow for intra-phagocyte survival and intracellular growth and proliferation to form hallmark granuloma structures [16]. This makes Arginase a potential target for therapeutic intervention during TB. In murine TB models, macrophage Arginase expression is associated with decreased bacterial killing and is an immunomodulatory target of *Mycobacterium tuberculosis* [16]. Similar observations have been described in fungal infections, with mice infected with Cryptococcus (*Cryptococcus gattii*) having elevated levels of Arginase expression in lung tissues [50]. In human-monocyte-derived macrophages, *Candida albicans* infection induces Arginase that blocks host NO production as a fungal survival mechanism via chitin exposure [41]. Our findings using the *arg2:GFP* line are consistent with these mammalian observations, with early *arginase* expression observed in innate immune cells after Mm and fungal infection. Further investigation is required to understand the molecular mechanisms of this intriguing host-pathogen interaction.

Arginase has been described as an anti-inflammatory macrophage marker due, in part, to high expression levels observed in mouse monocytes cultured with anti-inflammatory stimuli such as IL-4 or IL-13 to make M2/anti-inflammatory macrophages and observations in other murine macrophage models [51]. We observed macrophages expressing *arg2:GFP* during infection and at wound-healing stages of tailfin transection. Our findings complement studies on a zebrafish pro-inflammatory macrophage *tnfa* transgenic line, which suggest that an anti-inflammatory population of macrophages exist based on the switching off of *tnfa:GFP* in some macrophages during the wound-healing phase of tailfin transection, and are consistent with anti-inflammatory macrophages being present in the developing zebrafish larvae [45]. However in both inflammation and infection, *arginase*-expressing macrophages were much less frequent than *arginase*-expressing neutrophils, and neutrophil *arginase* expression predominated. It is important to note that the *mpeg1* promoter used to mark macrophages in our study is downregulated by Mm infection [52] therefore it is possible that our observations using the *mpeg1:mCherry* line is an underestimation of the population of macrophages that express *arg2:GFP* during Mm infection. This seems likely as there are many *arg2:GFP* positive granuloma-associated cells, some with phagocytosed bacteria, that expressed *arg2:GFP* but were neutrophil-marker negative, indicative of macrophages that lack visible *mpeg1:mCherry* marker. The granuloma-associated *arg2:GFP* observed may also be expressed in epithelioid-like cells that make up a large proportion of the zebrafish Mm granuloma structure, some of which are macrophage derived but may have lost macrophage markers [44]. Here, we have identified *mpeg1:mCherry* positive granuloma associated macrophages that express *arginase 2*, however it remains unclear as to how many macrophages are polarised towards this potential anti-inflammatory phenotype in zebrafish. Characterising the macrophage polarisation infection response fully will require further study for which the *arg2:GFP* zebrafish line will be an important tool.

Outside of immune cells, the *arg2:GFP* line has also illuminated arginase expression in the liver in highly infected individuals and in ionocytes, during both resting and inflammatory states. Arginase is a well-characterised liver enzyme [53,54] therefore expression here was not unexpected, though the function of the upregulated liver expression in highly infected individuals remains unclear. Interestingly, ionocytes have recently been described as a new airway epithelial cell type in human and mice and it is these cells that most highly express cystic fibrosis transmembrane conductance regulator (CFTR) the anion channel that is mutated in cystic fibrosis patients [55]. The *arg:GFP* line is a new tool that could be used to investigate the roles of these intriguing cells *in vivo*.

Our data indicate that *arginase 2*, an important anti-inflammatory mediator, is produced early after immune challenge, predominantly by neutrophils. The *arg2:GFP* line is an exciting addition to the zebrafish transgenic toolbox with which to investigate innate immunity during infections. It has the potential to be applied to multiple zebrafish disease models of infection and inflammation and may also be relevant to any models with an inflammatory component, from ageing to cancer.

## Acknowledgements

The authors would like to thank The BSU Aquarium Team for fish care and the IICD Technical Team for practical assistance (University of Sheffield). We would like to thank the Renshaw and Johnston groups for sharing their expertise in BAC transgenesis, especially Miss Catherine Loynes and Dr Stone Elworthy. Thanks to Dr Simon Johnston and Dr Stella Christou for invaluable help and expertise in fungal infection models. Thanks to Prof Stephen Renshaw for his helpful comments and discussions on the manuscript.

## Author Contributions

Conceived and designed the experiments: FRH, AHM, GFW, PME. Performed the experiments: FRH, AL, HEA, LGW, PME. Generation of *TgBAC(arg2:eGFP)sh571* transgenic zebrafish: AL. Analysed the data: FRH, AL, PME. Wrote and drafted the manuscript: FRH, AL, AHM, GFW, and PME.

