## Supplemental Figures and Legends for "An *arginase 2* promoter transgenic illuminates anti-inflammatory signalling in zebrafish" (from another version of this preprint: version 1, 2022-02-14)

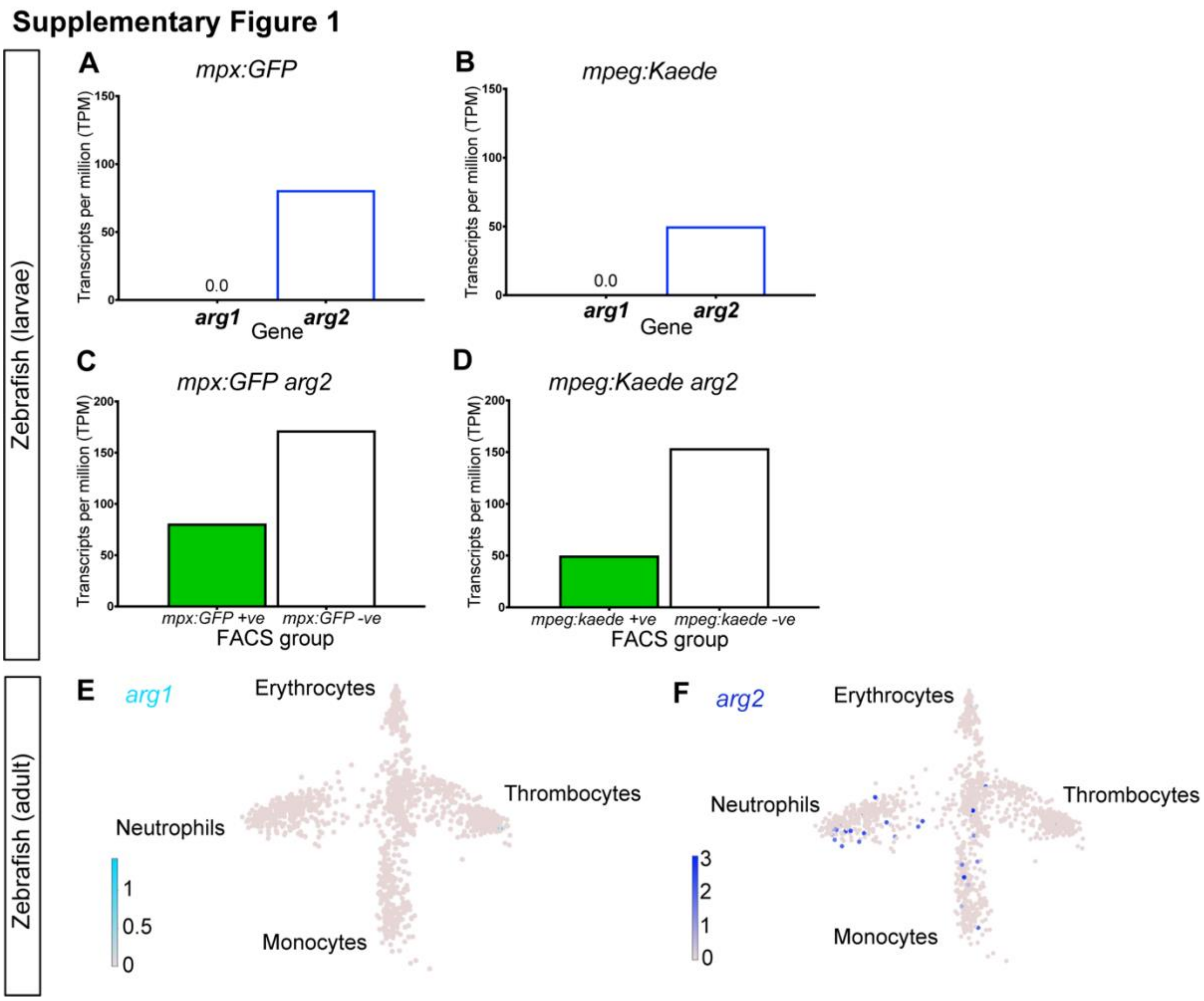


**Supplemental Figure 1: Zebrafish leukocytes express more *arg2* than *arg1* in both larvae and adults**

(A-D) Zebrafish neutrophils sorted via FACS of ~30 pooled 5dpf *Tg(mpx:GFP)i114* larvae and RNAseq performed on *mpx:GFP*+ (neutrophils, A and C) or *mpx:GFP*- (rest of fish) cells (C). Zebrafish macrophages sorted via FACS of ~30 pooled 5 days post fertilisation (dpf) *Tg(mpeg1:Gal4-VP16)gl24/(UAS-E1b:Kaede)s1999t* larvae and RNAseq performed on *mpeg:Kaede*+ (macrophages, B and D) or *mpeg:Kaede*- (rest of fish) cells (D). Datasets obtained from Rougeout *et al*., 2019. Data shown is an average value from 3 replicates.

(E-F) Gene expression of adult zebrafish leukocytes was determined using the zebrafish blood atlas (Athanasiadis *et al*., 2017, [37]). Each point represents separate scRNAseq replicates performed across multiple zebrafish wildtype and transgenic strains. Each arm of schematic indicates separate blood cell population (labelled). Deeper colour indicates higher expression (log10 scale bars described for each gene).

**Supplemental Figure 2**

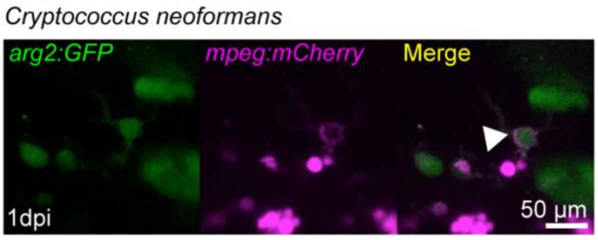


**Supplemental Figure 2: Macrophage *arg2:GFP* expression at 1dpi with *Cryptococcus neoformans***

Fluorescence confocal micrographs of *arg2:GFP* crossed to *mpeg:mCherry* after *Cryptococcus neoformans* infection at 1dpi with *arg2:GFP* positive macrophage indicated by arrowhead.

**Supplemental Figure 3**

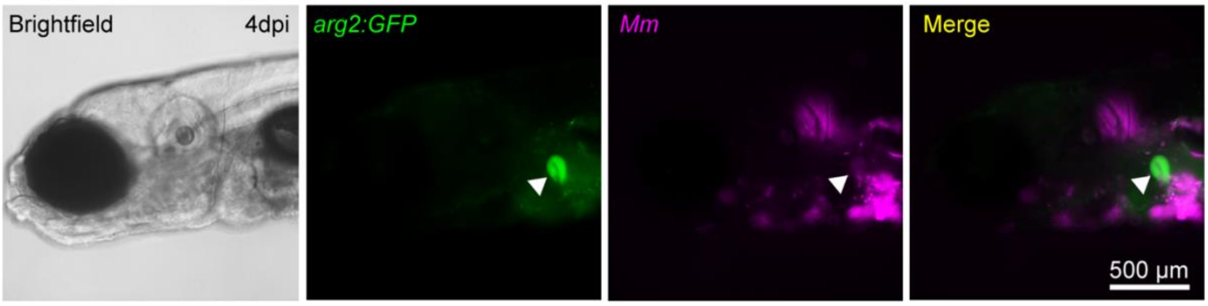


**Supplemental Figure 3: Mm infection upregulates *arg2:GFP* in the liver**

Brightfield and fluorescence micrographs of *TgBAC(arg2:GFP)sh571* larvae at 4dpi after Mm infection showing *arg2:GFP* liver expression with heavy levels of infection.
